# Artemis: Harnessing Knowledge Graphs for Next-Generation Drug Target Prioritization

**DOI:** 10.64898/2026.01.27.701959

**Authors:** Vladimir Yu. Kiselev, Edward Ainscow

## Abstract

Knowledge graphs (KGs) have become an important asset in biomedical research and drug discovery by enabling the structured integration of heterogeneous biological knowledge. When combined with machine learning (ML), KGs support the identification of novel drug-target relationships, but existing approaches are often KG-centric, relying primarily on graph structure and embeddings while overlooking disease-specific biological and clinical context. Moreover, many high-impact applications depend on proprietary KG infrastructures, limiting accessibility for the broader research community.

Here, we introduce Artemis, a practical and generalisable machine-learning framework for indication-aware target prioritisation that integrates public biomedical KGs with clinical evidence from the ChEMBL database. Artemis derives graph-based representations of clinically validated drug targets from multiple publicly available KGs and augments them with disease-relevant clinical features from ChEMBL. This hybrid feature space is used to train supervised ML models across seven disease indications, with performance assessed via cross-validation and guided parameter optimisation. The framework is further evaluated on emerging breast cancer targets reported at the San Antonio Breast Cancer Symposium 2024, demonstrating its ability to prioritise novel candidates.

Overall, this work demonstrates that publicly available KGs can be used for actionable, translational target discovery when coupled with clinical data. Artemis provides an accessible, scalable, and cost-efficient alternative to proprietary KG platforms. Thereby offering a practical solution for researchers seeking to prioritise therapeutic targets in real-world drug discovery settings.

**Key Points:**

1. KG applications can support the identification of novel drug–target relationships but rely primarily on graph structure while overlooking disease-specific biological and clinical context.
2. Artemis performs indication-aware target prioritisation that integrates public biomedical KGs with clinical evidence from the ChEMBL database.
3. Artemis is evaluated on emerging breast cancer targets reported at the San Antonio Breast Cancer Symposium 2024, demonstrating its ability to prioritise novel candidates.
4. Artemis provides an accessible, scalable, and cost-efficient alternative to proprietary KG platforms offering a practical solution for researchers seeking to prioritise therapeutic targets in real-world drug discovery settings.

## Introduction

Since their implementation by Google in 2012, knowledge graphs (KGs) have become a foundational technology across a wide range of research fields and applications [1].Click or tap here to enter text. By 2020, KGs began to gain traction in biomedical research, attracting significant attention from several major pharmaceutical companies. For example, AstraZeneca developed the Biological Insights Knowledge Graph [2]to support internal applications such as discovery ranking of CRISPR screen hits to aid novel target discovery. Novartis employed KG embeddings to recommend patient selection, design and end points for clinical trials [3].Click or tap here to enter text. GlaxoSmithKline (GSK) applied KGs to improve detection power in small-cohort genome-wide association studies (GWASs) [4], and Boehringer Ingelheim proposed a novel KG construction framework [5] later used for indication expansion and drug repurposing [6].

While these examples demonstrate the versatility and value of KGs in pharmaceutical drug discovery and development, they rely on proprietary platforms, inaccessible to the broader research community. A few commercial providers - such as [Clarivate](https://clarivate.com/life-sciences-healthcare/consulting-services/research-and-development-consulting/knowledge-graphs/) and [Qiagen](https://digitalinsights.qiagen.com/biomedical-knowledge-base/)- offer access to curated KGs, but at a considerable cost. However, in parallel, several publicly available biomedical KGs have emerged, beginning with the influential Hetionet KG [7]. AstraZeneca maintains a public [GitHub repository](https://github.com/AstraZeneca/awesome-drug-discovery-knowledge-graphs/blob/main/chapters/drug_discovery_kgs.md) cataloguing many of these open KGs. Community initiatives have also facilitated access and experimentation: for example, the PyKEEN framework [8] currently integrates nearly 40 standardized KGs, several of which are relevant to biomedical applications.

Recent studies on target discovery using KGs have largely focused on predicting potential drug targets by exploiting either improved graph construction, graph structural properties or by developing more advanced machine learning (ML) methods for KG representation and analysis [9–21].Click or tap here to enter text. While these approaches have led to methodological advances, they are fundamentally KG-centric: they treat the graph as the primary source of information and typically overlook disease-specific biological features. As a result, they do not explicitly incorporate the clinical or mechanistic context of the indication for which new targets are being sought.

ChEMBL is a large, open-access, expertly curated database of chemical structures linked to quantitative bioactivity measurements against defined biological targets [22].Click or tap here to enter text. Its standardized assays, target annotations, and chemical representations have made it a cornerstone dataset for training and benchmarking machine-learning models in drug discovery [23].Click or tap here to enter text. Despite its widespread use in ML, ChEMBL has rarely been integrated with biomedical KGs for target discovery, and systematic approaches that combine KG-derived features with clinical-stage evidence remain largely unexplored.

Here, we describe Artemis, a machine-learning framework designed to bridge network-level biological knowledge and clinical evidence for target evaluation. Artemis leverages publicly available biomedical KGs to represent known clinical drug targets and enhances these representations with disease-specific features derived from ChEMBL. This integrated feature space underpins supervised ML models trained across seven disease indications, enabling target evaluation that reflects both biological connectivity and empirical drug–target interactions. Model performance is validated using cross-validation and parameter optimisation and further tested on emerging breast cancer targets reported at the San Antonio Breast Cancer Symposium 2024 [24].

This approach addresses a limitation of existing KG-centric methods by moving beyond purely structural graph signals toward indication-aware target prioritization. Artemis is particularly suited for researchers who identify candidate targets from experimental [25] or computational screens and seek to rank them using an interpretable, ML-driven framework grounded in both KG topology and clinical evidence. By relying exclusively on publicly available resources, Artemis provides a novel, practical, and accessible strategy for target evaluation with direct relevance to real-world drug discovery workflows.

## Methods

### Knowledge Graphs

Every KG consists of nodes (entities, such as genes, proteins, drugs, or diseases) and edges (relationships that capture known or inferred biological interactions or associations, derived from curated databases, experimental evidence, or computational predictions). Two nodes connected by an edge create a triple. Triples are building blocks of a KG. In our study, we focused on several publicly available biomedical KGs, all of which are included and standardized within the PyKEEN python library [8].Click or tap here to enter text. Their properties are shown in Table 1.

**Table 1.**
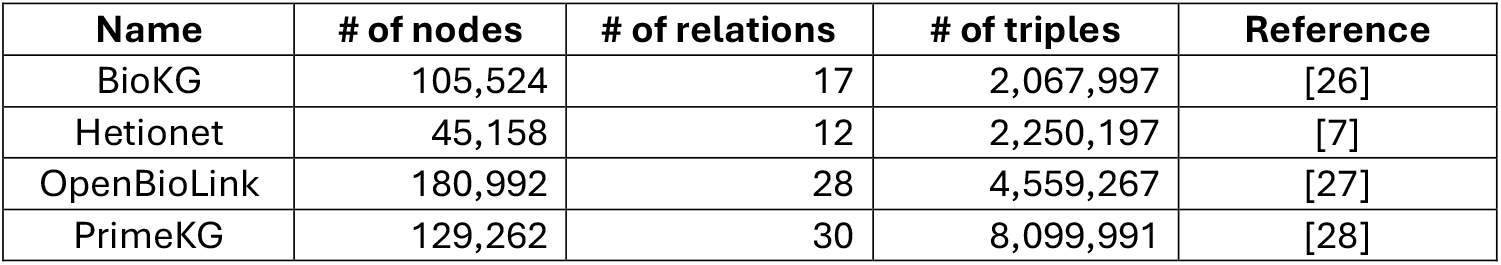
Characteristics of KGs used in the study.

These KGs were selected to provide a diverse and representative set of biomedical networks varying in scale, ontology structure, and data provenance, ensuring a robust evaluation of our target discovery framework. Full details of the KGs are available in the Supplementary Notebook.

### Knowledge Graph Embeddings

Knowledge graph embedding (KGE) is a ML technique that refers to the process of learning low-dimensional vector representations of entities and relations in a KG while preserving its structural and semantic characteristics.

We generated embeddings of the KGs using the PyKEEN framework, that provides standardized implementations of a wide range of KGE algorithms and offers automated hyperparameter optimization, dataset handling, and evaluation utilities.

For BioKG and Hetionet KGs, we adopted the model (RotatE) and best performing hyperparameter configurations reported in a recent study [29].Click or tap here to enter text. These configurations have been empirically validated to produce high-quality embeddings for biomedical relation prediction tasks. For the remaining KGs, we applied the same hyperparameters as used for Hetionet to maintain methodological consistency across datasets (due to a large graph size the *number of epochs* parameter for PrimeKG was reduced to 150). All triples of each graph were used for the embeddings.

### Knowledge Graph Link Predictions

Link prediction in KGs aims to infer previously unobserved relations between entities by leveraging the existing network structure and the learned embeddings of entities and relations. In addition to discovering new associations, link prediction can also be used to quantify the confidence or plausibility of existing relations by assigning a score to each known triple within the KG.

For each KG, we performed link prediction using the PyKEEN framework. Rather than predicting all possible links across the entire graph, we adopted a gene-centric approach, focusing exclusively on links between gene/protein nodes and other relevant entities, depending on the relation types defined within each KG. Because relation naming conventions vary across public KGs, we manually curated a subset of relations corresponding specifically to gene and protein associations. A detailed list and description of these curated relations are provided in the Supplementary Notebook.

The output of the link prediction process was a gene–node matrix, where rows represent genes/proteins, columns correspond to other KG entities, and cell values denote the predicted strength of the relation between each pair. This matrix includes only the relation types already present in the original KG, ensuring that predictions remain biologically interpretable and consistent with the underlying graph schema.

### Clinical Scores

To augment the KG-derived relationships with clinical evidence, we queried ChEMBL database (version 36) for bioactive drug molecules associated with clinical trial data and annotated therapeutic indications. All molecular variants (e.g. salts, formulations, and prodrugs) were standardised to their corresponding parent molecules, and associated information - including clinical trial phase, [Medical Subject Headings (MeSH) disease annotations](https://www.nlm.nih.gov/mesh/meshhome.html), mechanisms of action, and molecular targets - was aggregated at the parent-molecule level to generate a unified dataset. Disease indications were grouped by selecting a top-level MeSH branch and including all its descendant terms. This grouping strategy was applied to a set of cancer and non-cancer indications selected to reflect a combination of high disease prevalence, clinical impact, and availability of well-characterised therapeutic targets. The cancer indications represent some of the most common and intensively studied tumours, spanning distinct tissues of origin, molecular drivers, and therapeutic landscapes. The non-cancer indications were included to evaluate the generalisability of the framework beyond oncology, particularly in chronic, multifactorial diseases with extensive clinical and genetic evidence bases. Together, this selection enables a systematic assessment of model performance across diverse disease areas:

#### Cancer indications

- Breast Neoplasms (breast)
- Lung Neoplasms (lung)
- Intestinal Neoplasms (bowel)
- Prostatic Neoplasms (prostate)
- Skin Neoplasms (melanoma)

#### Non-cancer indications

- Diabetes Mellitus, Type 2 (diabetes)
- Cardiovascular diseases (cardiovascular)

To mitigate bias introduced by drugs which may act through multiple, closely related, proteins, we addressed target redundancy at the gene family level. For each drug– disease combination, we identified over-represented HGNC gene families [30] - defined as cases in which more than 40% of family members were targeted by a single molecule and retained a single representative gene per family (selected alphabetically).

To represent the clinical maturity of each target, we computed a clinical score based on the number and clinical phase distribution of associated clinical trials (we only considered trials from the last six years). Specifically, if a target was linked to ***a*** phase I trials, ***b*** phase II trials, ***c*** phase III trials, its score was calculated as:

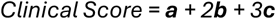

Additional approval bonus (+20) was added to the clinical score if a target has at least one approved drug in each disease indication. This weighting scheme assigns higher importance to targets evaluated in more advanced clinical stages, and even higher importance to the ones that are targeted by approved drugs, reflecting their increased translational validation and clinical utility.

The resulting clinical target scores were subsequently used for training the ML models described below. We considered three distinct training scenarios:

- (*All*) All clinical targets - including all targets associated with any clinical trial phase.
- (*Approved*) Approved targets - restricted to those with clinical phase value of 4 in ChEMBL for a given indication.
- (*Unique*) Indication-specific and shared targets - where only unique indication-specific targets or targets present across all indications were retained.

Supplementary Fig. 1 summarizes the number of clinical targets used in each scenario, showing both the per-indication counts and the overlap between disease areas.

### Model Training

We employed a supervised learning approach to train multiple ML models, using KG link prediction scores as input features and clinical scores as output labels. Because the clinical scores were continuous, we evaluated several modelling paradigms to explore both regression and classification performance:

- Regression models: linear regression and K-neighbour regression
- Multiclass classification models: Random Forest (RF) and Support Vector Machine (SVM)
- Binary classification models: Random Forest (RF) and Support Vector Machine (SVM)

Prior to training, all clinical scores were log_2_-transformed to reduce skewness and stabilize variance. To improve model robustness and prevent overfitting, we augmented the dataset with negative samples, defined as genes not present among the clinical drug targets. These “negative targets” were randomly selected from the human genome excluding known drug targets and assigned a clinical score of -1. For multiclass classification models, continuous clinical scores were discretized into quantile-based bins (3, 4 and 6 bins – see Fig. 1) using the *qcut()* function from the pandas python library. For binary classification models positive clinical scores were considered as ones and negative scores as zeros.

**Figure 1.**
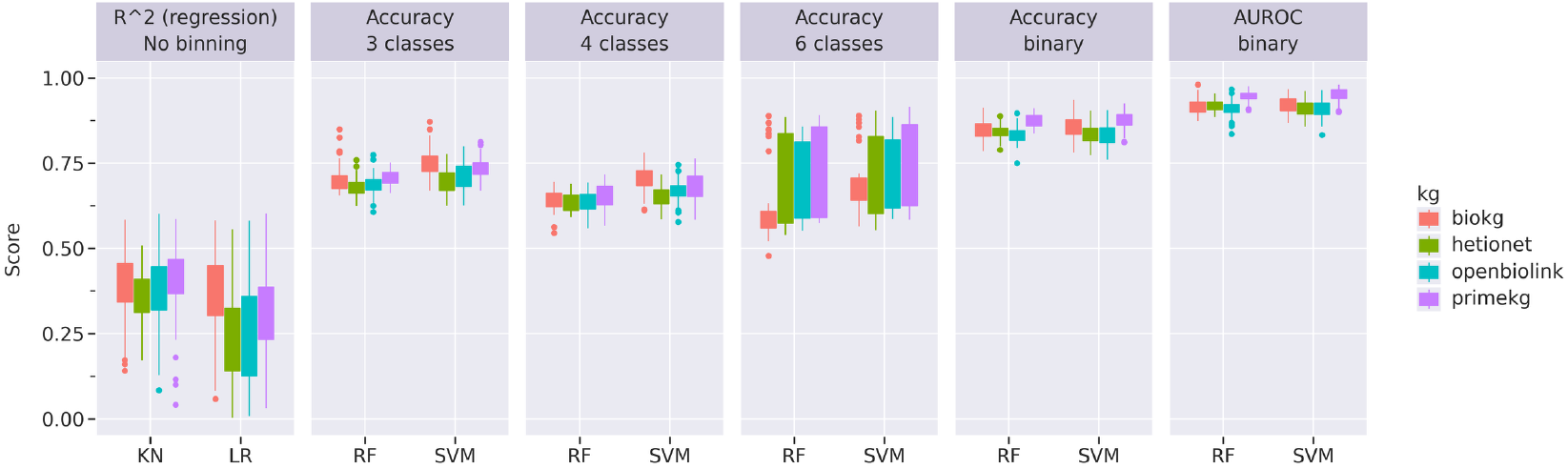
Cross-validation results of various regressors/classifiers. Each bar plot consists of 70 data points corresponding to 7 disease indications and 10 different random seeds for each indication. KN – k-neighbour regression, LR – linear regression, RF – random forest, SVM – support vector machines. R^2 - coefficient of determination, ARI – adjusted Rand index, AUROC – area under the ROC curve. The baseline metrics for each indication are shown in Supplementary Fig. 4.

### San Antonio Symposium Breast Cancer Targets

As an external validation set, we curated a list of emerging breast cancer targets from abstracts presented at the San Antonio Breast Cancer Symposium (SABCS) 2024 [24]. Abstracts were processed using a text-mining pipeline to identify sentences containing pharmacologically relevant keywords, including target(ing), inhibit(ing), degrade(r), agonist, directed, and bind(ing). Identified sentences were then screened for mentions of human protein-coding genes and their aliases, based on annotations obtained from the HUGO Gene Nomenclature Committee (HGNC) [30].Click or tap here to enter text. Candidate genes were cross-referenced against the set of known clinical breast cancer targets described above, and genes not present in this reference set were retained as putative novel therapeutic targets for breast cancer.

It is important to note that this external validation set was derived exclusively through text mining of the conference abstracts and therefore captures only the likely presence of a pharmacological drug-gene-disease association. As such we do not distinguish between those that provide a positive effect on the disease or potential off-target or negative impacts. In addition, all candidate genes identified through this process were treated equivalently, without applying any weighting to reflect the strength, frequency, or experimental maturity of the supporting evidence. Therefore, from this gene set we expect an enrichment in novel target genes, but that not all genes identified in this way to be potential drug targets for breast cancer.

### Pathway Genes

When working with SABCS to enhance the predictive performance and biological relevance of our models, we supplemented the training data for each indication with pathway-associated genes obtained using the Geneshot search engine [31]. The search terms used in pathway genes queries:

#### Cancer indications

- Breast Cancer (breast)
- Lung Cancer (lung)
- Bowel Cancer (bowel)
- Prostate Cancer (prostate)
- Melanoma (melanoma)

#### Non-cancer indications

- Diabetes Mellitus Type 2 (diabetes)
- Cardiovascular Disease (cardiovascular)

These genes, identified based on literature co-occurrence and functional enrichment, were assigned a positive clinical score of 1 and included alongside the clinical drug targets during model training.

The returned pathway genes were sorted by their rank and only the top *N* genes were used. Furthermore, only unique pathway genes - those not shared across multiple indications -and genes present in every indication were retained from the top *N* genes. This filtering step aimed to reduce redundancy and emphasize disease-specific molecular context.

Given that many clinical targets share common properties across indications, we hypothesized that incorporating pathway genes would provide additional indication-level specificity and improve the model’s ability to distinguish between disease contexts.

In the Results section we consider three scenarios corresponding to obtaining top 0, 100 and 300 pathway genes from the Geneshot search engine. These numbers change after the uniquification (Supplementary Fig. 2); however, we still refer to them as 0, 100 and 300 in the text.

## Results

### Cross Validation

#### Number of KG nodes

Given that there is considerable variation in size of the public domain KGs tested, we first assessed whether model performance was influenced by the size of the KG. To evaluate this, we conducted a cross-validation experiment using the Hetionet KG and breast clinical targets. The KG feature space - defined as the number of node embeddings used as model input - was systematically down sampled to subsets containing 30,000; 20,000; 10,000; 5,000; 1,000; 500; 100; 50; 10; and 5 features. The corresponding performance metrics across these feature subsets are shown in Supplementary Fig. 3.

Model performance remained largely stable across a broad range of feature sizes, with a notable decline only when the number of features dropped below approximately 1,000. To balance computational efficiency with predictive power for larger KGs, we therefore randomly selected 10,000 features for all KGs - one order of magnitude above the observed performance threshold - for link prediction and subsequent model training.

#### Performance

To assess the predictive performance of the ML models, we conducted five-fold cross-validation for each regressor and classifier (Fig. 1). For the multiclass classification experiments, we tested models with 2, 3, 4, and 6 discrete classes derived from the binned clinical target scores.

Across all experiments, the Random Forest (RF) and Support Vector Machine (SVM) models consistently achieved the highest predictive performance, particularly in the binary classification setup. Based on these results, subsequent analyses and target-ranking experiments in this study were performed using the RF model trained on binary clinical scores.

### Model Performance

To evaluate model performance, we assessed several metrics, focusing initially on specificity and sensitivity. These metrics were quantified by measuring the overlap between predicted targets and the corresponding clinical target sets for each disease indication. Overlaps were visualised as heatmaps, with disease indications represented on both axes (Fig. 2, Hetionet example). When models were trained using *All* clinical targets (see Methods) and evaluated with the default random forest (RF) probability threshold (Fig. 2, left), the resulting heatmap was largely homogeneous, with values close to 100% across indications, showing limited specificity. In contrast, applying a stricter RF threshold and restricting the training set to *Unique* clinical targets (Methods, Fig. 2, right) substantially reduced off-diagonal overlap, while diagonal values remained close to 100%. This pattern indicates that the models retained the ability to recover their respective training targets while achieving improved indication specificity.

**Figure 2.**
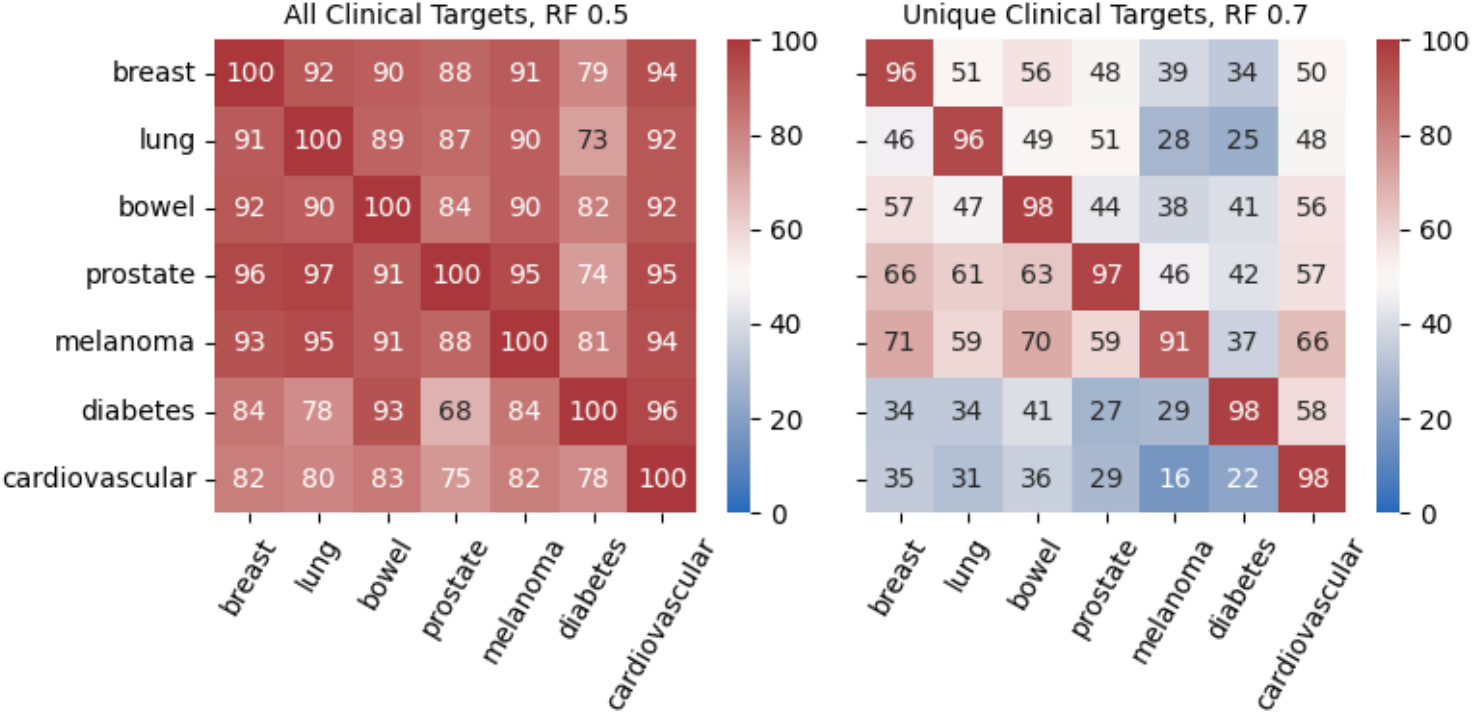
Heatmaps showing percentages of common genes between predicted targets from x-axis indications and the input clinical targets (y axis) for Hetionet KG. Left subfigure corresponds to models training on *All* clinical targets with 0.5 RF probability threshold. Right subfigure corresponds to models training on *Unique* clinical targets with 0.7 RF probability threshold. Each cell is an average of 10 independent results corresponding to 10 different random seeds. Default RF probability threshold of 0.5 was used for model predictions. The results for the other KGs and other RF thresholds (0.6-0.9) are shown in Supplementary Figs. 5-9.

### Based on these observations we introduced the following custom metrics

- Specificity was defined as one minus the average proportion of false positives across all other indications (i.e., one minus the mean of the off-diagonal values divided by 100).
- Sensitivity was defined as the proportion of true positives for each indication (i.e., the diagonal value divided by 100).

Intuitively, higher specificity corresponds to fewer off-target predictions (lower off-diagonal elements), while higher sensitivity indicates stronger recovery of known clinical targets (higher diagonal elements).

In addition to specificity and sensitivity, we introduced an additional performance metric to capture the overall predictivity of the model. Specifically, the baseline prediction level was defined as the fraction of all genome genes that are predicted as targets for a given indication. This metric provides a global measure of model stringency, independent of any disease context. Baseline predictivity is particularly informative for target prioritisation tasks, where the practical utility of a model depends not only on its ability to recover known targets but also on its capacity to restrict predictions to a manageable and biologically meaningful subset of genes. A high baseline prediction level indicates a permissive model that nominates a large fraction of the genome as potential targets, reducing prioritisation value and increasing experimental burden. Conversely, a low baseline reflects a more selective model, yielding shorter, more actionable target lists suitable for downstream validation.

By jointly considering baseline predictivity alongside sensitivity and specificity, we ensure that model performance balances biological coverage, indication relevance, and practical usability in real-world drug discovery workflows.

### Parameter Optimization

Our pipeline incorporated several tuneable parameters, including the probability threshold used for Random Forest (RF) predictions, as well as the number of clinical targets included during model training. To identify the optimal parameter configuration, we performed a parameter optimization analysis using the three previously defined metrics – baseline, specificity and sensitivity. Optimization was conducted across the following parameter space:

- Clinical targets: *All, Approved, Unique* (see Methods)
- Random Forest probability threshold: *0*.*5 (default), 0*.*6, 0*.*7, 0*.*8, 0*.*9*

The objective of the optimisation was to jointly maximise specificity, sensitivity, and baseline performance. The analysis revealed several consistent trends (Fig. 3). As expected, specificity increased with more stringent filtering, both when progressively restricting the clinical target set *(All → Approved → Unique)* and when increasing the RF probability threshold *(0*.*5 → 0*.*9)*. Sensitivity exhibited the opposite trend, although the decrease was comparatively modest. In contrast, baseline performance declined sharply with increasing RF probability thresholds but was consistently higher when the clinical target set was more restricted. Based on these trade-offs, the optimal parameter range was defined as:

**Figure 3.**
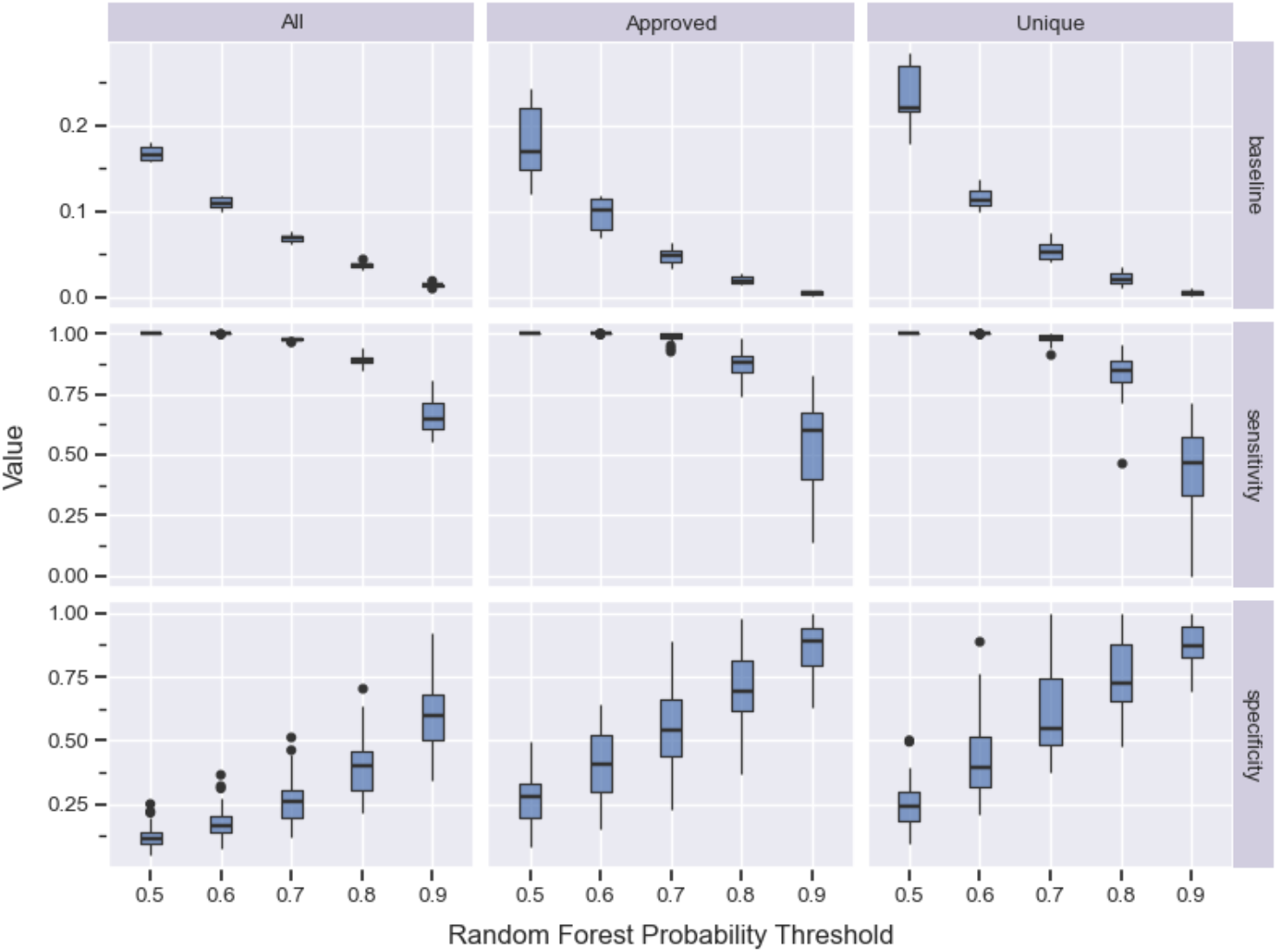
Optimisation of specificity sensitivity and baseline metrics. Boxplots correspond to all KGs and all disease indications.

- Using *Unique* clinical targets, and
- Setting the RF probability threshold to *0*.*7*.

### San Antonio Symposium Breast Cancer Targets

Having established the optimal parameter range, we next evaluated Artemis ability to identify drug targets beyond those already known from clinical data. As an external validation set, we used emerging novel breast cancer targets extracted from the San Antonio Breast Cancer Symposium (SABCS) 2024 abstracts (Methods).

In the same manner as in Fig. 2 we calculated the overlap between the predicted targets and the SABC targets for each indication and each KG (Fig. 4). Note that we expected that the text mining approach to identify the novel targets would contain a degree of inaccuracy in identifying gene names within an abstract to being a legitimate investigational drug target, therefore 100% prediction would not be expected.

**Figure 4.**
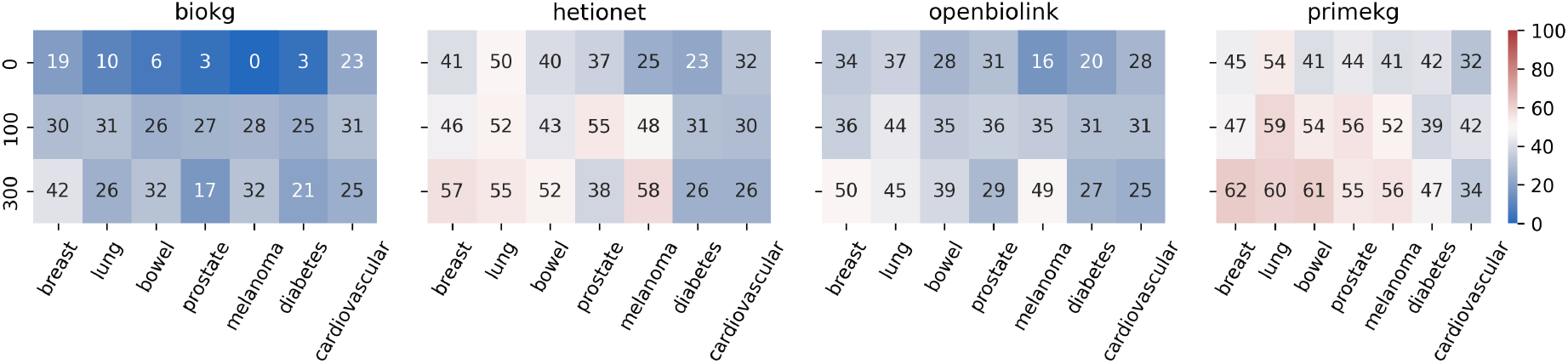
A heatmap showing percentages of common genes between predicted targets from x-axis indications and the SABCS novel targets. Y-axis represents the number of pathway genes used for training (Methods). Each cell is an average of 10 independent results corresponding to 10 different random seeds. The results for the non-default RF thresholds are shown in Supplementary Figs. 10-14.

Although the breast cancer model was not consistently the top-performing model, a clear separation emerged between cancer and non-cancer indications. Models trained on non-cancer diseases (diabetes and cardiovascular disease) showed minimal overlap with targets reported at SABCS, whereas models trained on cancer indications exhibited prediction patterns like the breast cancer model. This convergence suggests that shared oncogenic mechanisms are captured by the KG-derived features and contribute to cross-cancer target prioritisation.

Incorporating pathway genes into the training set (Methods), using either 100 or 300 genes, consistently increased the number of predicted targets, indicating that pathway-level information provides complementary signal beyond clinically validated targets alone. This effect was substantially more pronounced for cancer indications than for non-cancer diseases, consistent with the notion that cancers share common oncogenic pathways that are effectively captured by the added pathway context. Notably, in several settings, inclusion of pathway genes led to the strongest overall performance for the breast cancer model, showing the desired outcome.

Importantly, the positive classification rate for novel SABCS targets was substantially higher than the baseline prediction level (6–7%; Fig. 3), indicating that the models’ predictions are markedly enriched relative to random expectation.

Furthermore, we compared the RF probability scores assigned to SABCS genes across different KGs (Fig. 5). Genes were grouped using hierarchical clustering, revealing three major clusters corresponding to high, intermediate, and low RF probability levels. Overall, the results indicate strong consistency in predictions across KGs. Where discrepancies between KGs were observed - primarily within the intermediate cluster - RF probabilities for ∼60-70% of genes clustered around 0.5, suggesting uncertainty rather than systematic disagreement. In contrast, the remaining ∼30% of genes showed larger divergences in RF probabilities, suggesting KG-specific effects.

**Figure 5.**
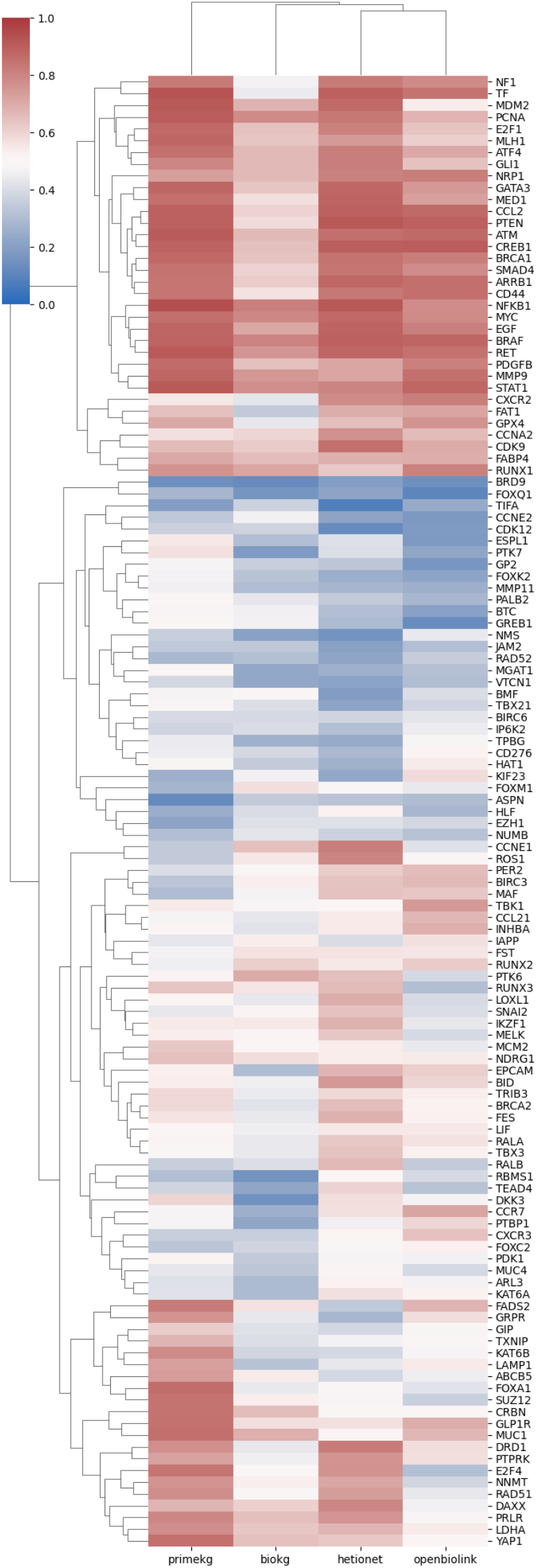
A heatmap of predicted RF probabilities for SABCS novel breast cancer targets from a breast model without pathway genes (Methods). Cell values correspond to a mean across 10 runs with different random seeds. Only genes that were present in all four KGs are present in the heatmap (genes that were not present in at least one KG were filtered out). Genes and KGs were clustered by hierarchical clustering that are shown as the dendrograms.

The top-ranked predicted genes included several well-established players in breast cancer biology, many of which remain undrugged despite extensive evidence supporting their roles in tumorigenesis. These included BRCA1, PTEN, and MED1 among others. FAT1 is target of extensive current research due to its association with the emergence of resistance to CDK4/6 inhibitors [32] and its role in the Hippo/YAP pathway. Other targets in this bracket are those that have been explored for other oncology indications (e.g. BRAF, MDM2 and CDK9) and therefore could be considered as potential areas for drug repurposing or for indication expansion. The analysis also selected genes such as GATA3 which is a differentiation marker of luminal breast cancer and associated with positive prognosis [33]. Indicating that activation of such a gene (or related pathway) could be a therapeutic strategy.

The mid-ranked portion of the predictions was particularly intriguing, as it highlighted divergence between KGs and may represent the most novel target space. Within this group, we observed several emerging targets such as YAP1 and TEAD4, key mediators of contact inhibition pathways that are frequently dysregulated in cancer. Their presence, alongside FAT1 in the top-ranked cluster, substantiates the potential of the Hippo/YAP pathway as therapeutic targets. Additionally, RAD51 overexpression has been associated with resistance to PARP inhibitors, while RUNX2 and RUNX3 have been associated with aggressive, metastatic forms of breast cancer through their action in stabilising MYC [34].

The lower-ranked genes presented a more heterogeneous mix. Some, such as CDK12 and RAD52, are likely false negatives, as they are well-established in DNA repair pathways and PARP inhibitor resistance. CD276 (B7-H3) a surface antigen which has been reported to be overexpressed in many cancers, but antibodies and CAR-T therapeutics against this target have had limited clinical success to date. The remaining genes are less characterized, representing potentially novel candidates for further biological validation.

## Discussion

In this work, we introduced a practical framework Artemis that harnesses several publicly available KGs to systematically identify and prioritise therapeutic targets for defined disease indications. The approach aggregates network-derived characteristics of already established drug targets, uses these features to train an ML model, and then applies the model to score previously uncharacterised candidates. Importantly, the training set is anchored in targets supported by clinical evidence of efficacy within the relevant disease context. We demonstrated that this methodology is broadly applicable by constructing models for a range of cancer and non-cancer indications and observing consistently strong predictive performance. Overall, this study showed that publicly accessible KGs in combination with the clinical data can be used for actionable drug discovery, offering a practical and resource-efficient alternative to bespoke proprietary KG systems.

We assessed Artemis’ performance using cross-validation and introduced quantitative metrics to guide parameter optimisation. This procedure identified a RF model with a probability threshold of 0.7 as the best-performing configuration.

Our results also indicated that the choice of KG had only a modest impact on overall performance. This suggested that Artemis is flexible and can be readily extended to incorporate newly updated or more sophisticated public KGs as they become available.

We further evaluated Artemis on a set of emerging breast cancer targets extracted from San Antonio Breast Cancer Symposium (SABCS) 2024 abstracts (Methods). For all KGs the breast cancer prediction levels were significantly higher than a baseline level. The results also showed similarities in cancer indication predictions suggesting shared oncogenic mechanisms captured by the KG-based features. Even though the prediction levels of non-cancer indications were lower than that of cancer ones, they were still above the baseline levels. While adding pathway genes to the training set did not substantially alter performance on known clinical targets (Fig. 3), it had a pronounced impact on the prediction of novel candidates from SABCS list (Fig. 4), underscoring the value of pathway-derived features for identifying previously uncharacterised disease-relevant genes.

Examination of the highest-ranked SABCS predictions revealed several well-established genes in breast cancer biology, many of which remain undrugged despite extensive evidence supporting their relevance. At the opposite end of the ranking, the weakest predictions were largely genes with limited or no known involvement in breast cancer, although a small number were likely false positives. The mid-ranked group was particularly noteworthy, as it contained genes that may represent genuinely novel target candidates.

These observations suggest that relying solely on KG-derived features even with the clinical data may still be insufficient for fully resolving target novelty or prioritisation. Incorporating additional, orthogonal ranking sources from publicly available resources could further refine prediction quality. For example, a recent approach [35] integrates multiple independent ranking signals and applies an optimisation algorithm to identify the most promising targets - a strategy that could complement and enhance Artemis’ prioritisation pipeline.

This work illustrates the value of applying clinically grounded, knowledge graph–driven machine learning to therapeutic target prioritisation. This is particularly useful where modern genomic approaches, such as CRISPR screening, single cell expression analysis and sequencing of patient sample banks, generate lists of 10-100s of genes as potential therapeutic targets. While the analyses presented here focus on selected cancer and non-cancer indications, the underlying methodology is disease-agnostic and make Artemis readily transferable to other disease areas where there is a critical unmet need for novel therapeutic targets.

## Supporting information

Supplementary Notebook

## Acknowledgements

The authors would like to thank Bethan Psalia and Rohit Batta for their insightful feedback.

## Author contributions

Vladimir Yu. Kiselev (Conceptualization, Methodology, Investigation, Software, Writing-original draft, Writing - review & editing) and Edward Ainscow (Conceptualization, Supervision, Writing - review & editing)

## Data and Code Availability

All external and generated data used in this study are publicly available in the Amazon Web Services (AWS) S3 bucket (*s3://alethiotx-artemis*), which can also be browsed at http://alethiotx-artemis.s3-website.eu-west-2.amazonaws.com/.

The core analyses are implemented in Python and provided in the *artemis* module of the *alethiotx* Python package (https://pypi.org/project/alethiotx/), with full documentation available at https://alethiotx.github.io/pypi/alethiotx/artemis.html.

A fully reproducible Nextflow [36] pipeline for generating all data and figures in this manuscript is available at https://github.com/alethiotx/artemis-paper. Additional Nextflow pipelines for KG embedding generation and KG link prediction are available at https://github.com/alethiotx/artemis-kgs-embeddings and https://github.com/alethiotx/artemis-kgs-link-predictions, respectively. All workflows were executed on the Seqera Platform using AWS Batch and read from and write to the same S3 bucket; embedding generation leveraged p3 instances with NVIDIA V100 GPUs.

### Use of Artificial Intelligence (AI) Tools

AI-based language models were used solely to assist with formatting of text and drafting of limited code examples. No AI tools were used to generate scientific conclusions, results, or interpretations. All outputs were reviewed and validated by the authors, who retain full responsibility for the content of this manuscript.

