## Supplementary Figures for "Artemis: Harnessing Knowledge Graphs for Next-Generation Drug Target Prioritization"

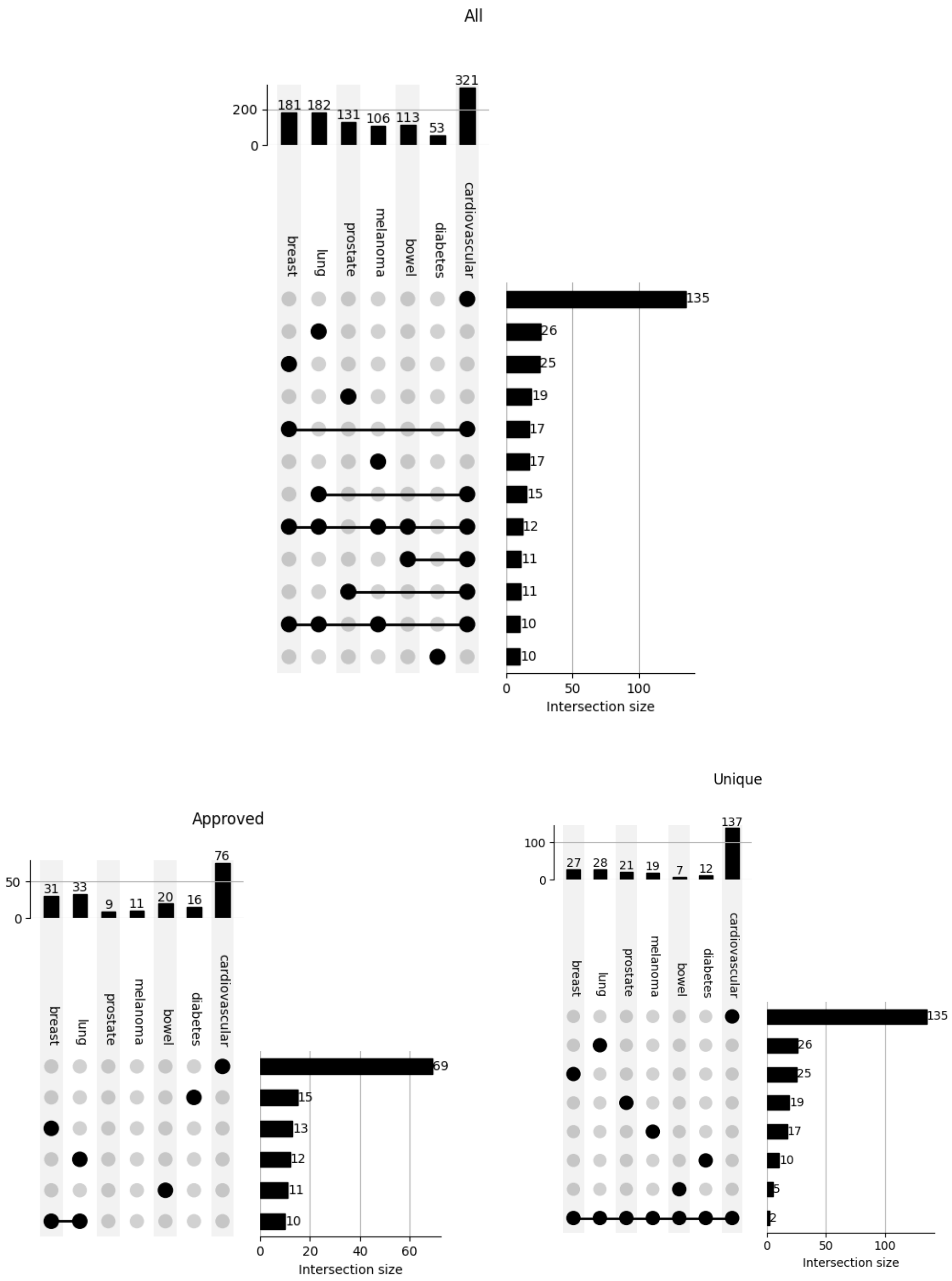


Figure 1. Upset plots (https://upsetplot.readthedocs.io) showing the number of clinical targets (obtained on 2025-12-12) used in model training after selecting either: 1. All the targets (top subplot), not showing intersections with the size smaller than 10; 2. Only FDA approved targets (bottom left subplot), not showing intersections with the size smaller than 10; 3. Either unique indication-specific targets or targets present in all indications (bottom right subplot).

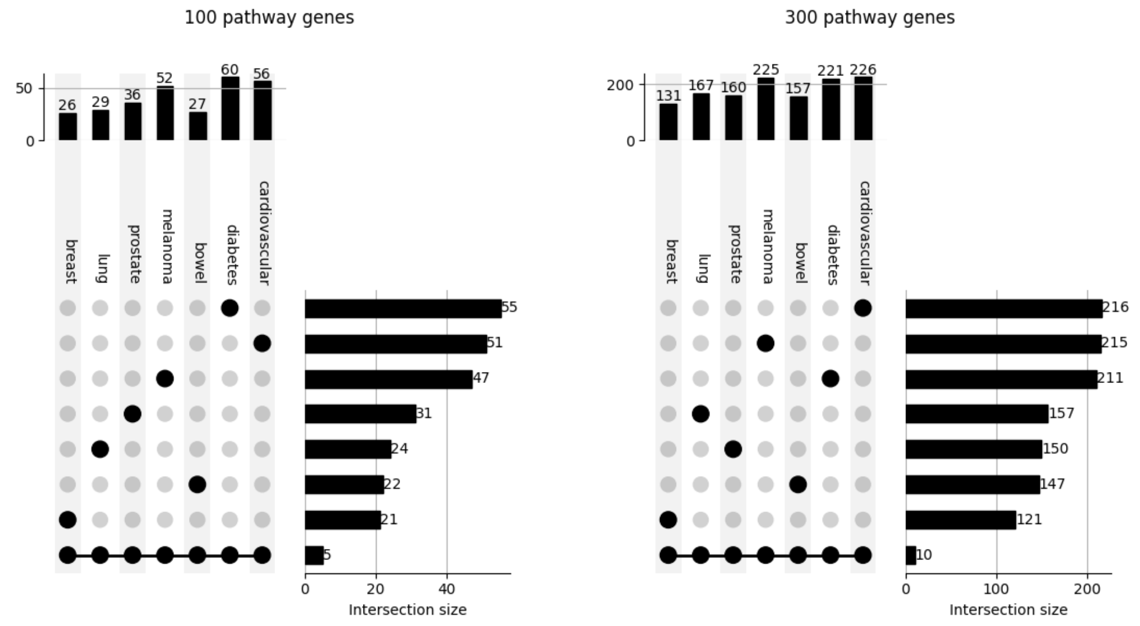


Figure 2. Upset plots (https://upsetplot.readthedocs.io) showing the number of pathway genes (obtained on 2025-12-12) used in training after obtaining the top 100 (left) and 300 (right) pathway genes for each indication via Geneshot and selecting only the unique indication-specific genes and the genes that are present in all indications.

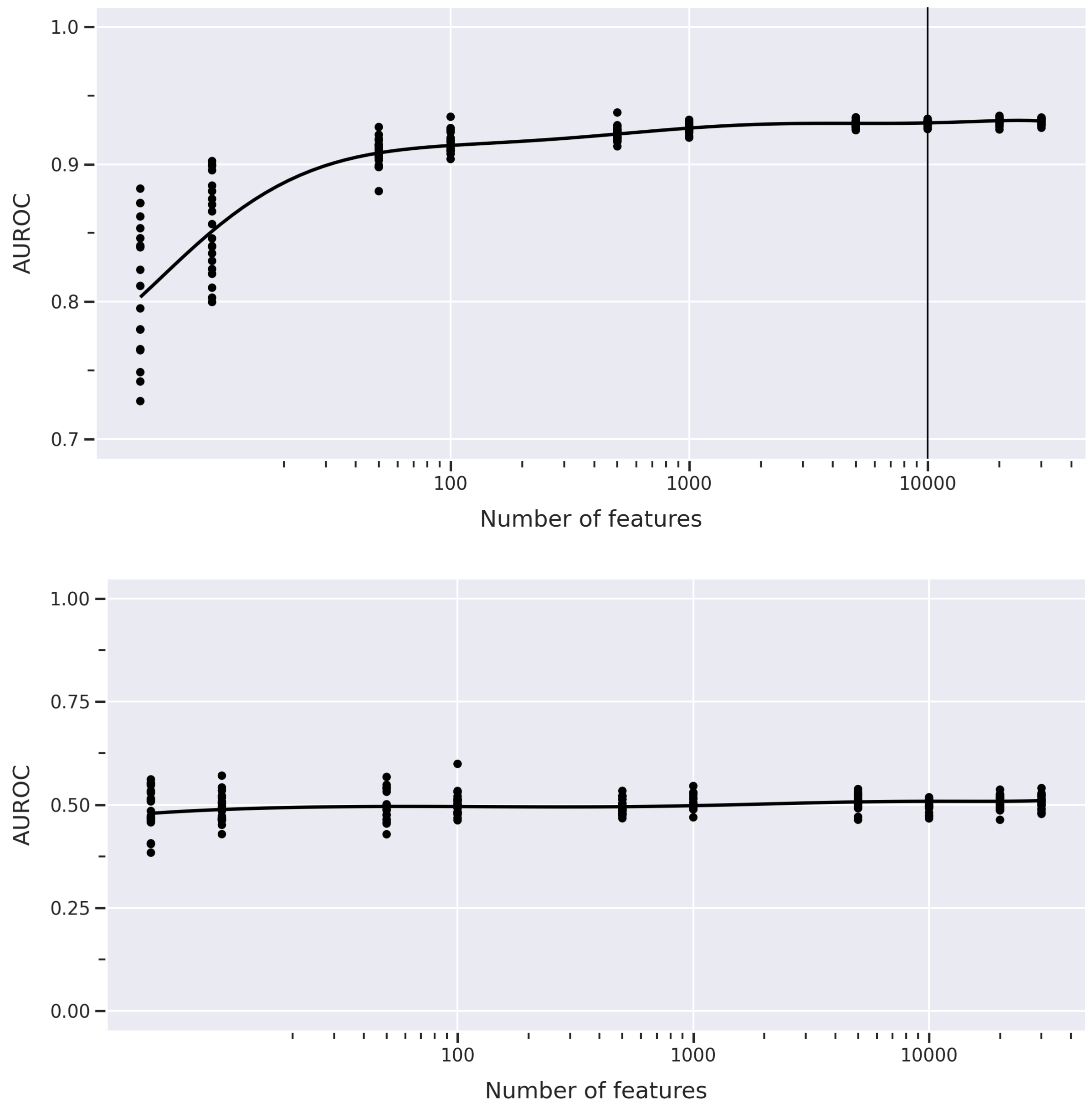


Figure 3. Top - cross-validation (5-fold) results of the random forest classifier based on the Hetionet KG and binary breast cancer targets. Bottom – same as the top figure but with the randomly shuffled training labels. For each point on the x axis KG features were randomly selected 10 times. Solid line corresponds to the polynomial fit with 7 degrees. Black vertical line corresponds to the threshold of 10,000 random features that was used in all KGs to select features for training.

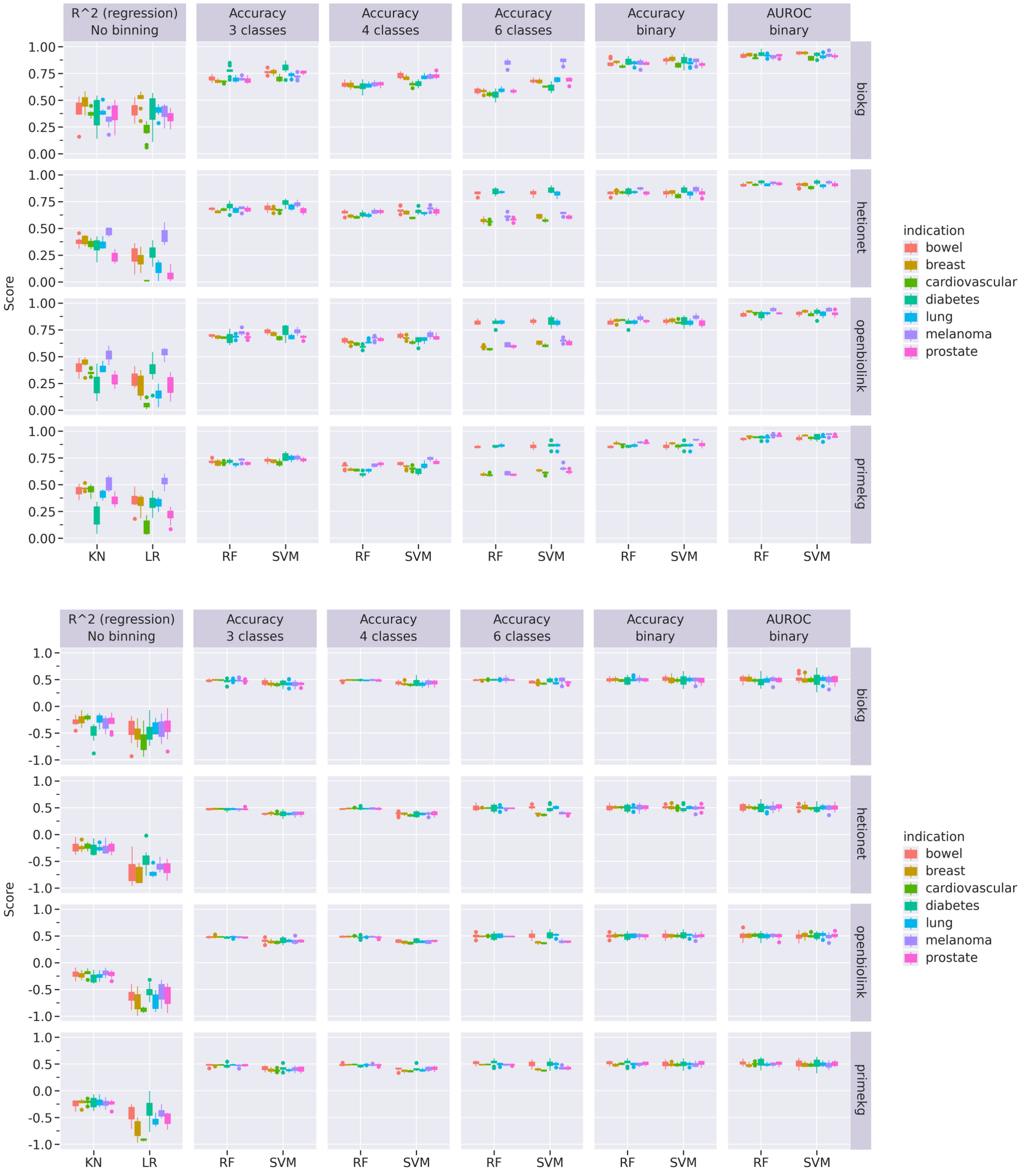


Figure 4. Cross-validation results of various regressors/classifiers. Top plot corresponds to real training targets; bottom plot corresponds to baseline performance when training target scores are randomly shuffled. Each bar plot consists of 10 data points corresponding to 10 different random seeds for each indication. KN – k-neighbour regression, LR – linear regression, RF – random forest, SVM – support vector machines. R^2 - coefficient of determination, ARI – adjusted Rand index, AUROC – area under the ROC curve.

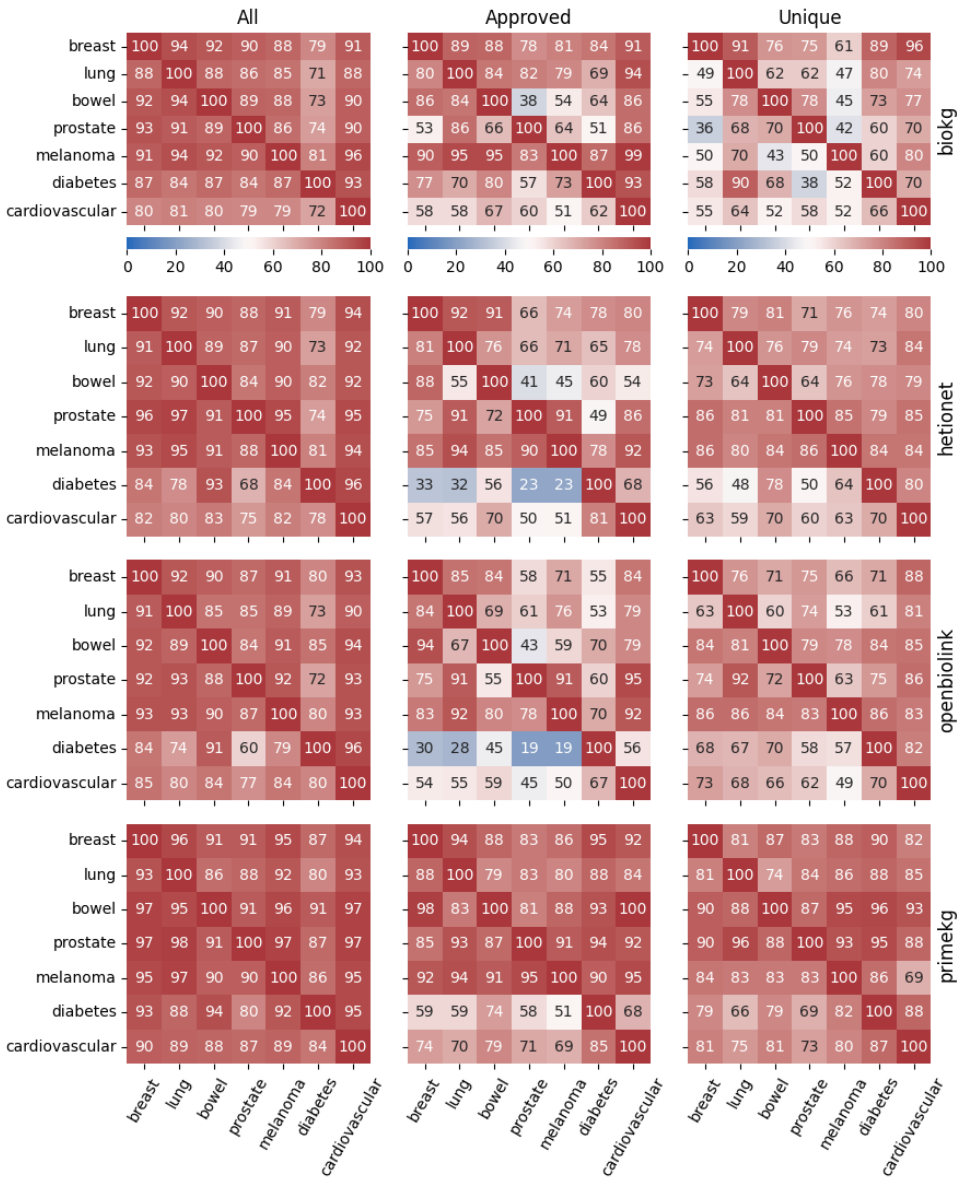


Figure 5. A heatmap showing percentages of common genes between predicted targets from x-axis indications and the input clinical targets (y axis). Each cell is an average of 10 independent results corresponding to 10 different random seeds. No pathway genes were used in training. RF probability threshold of 0.5 was used for model predictions.

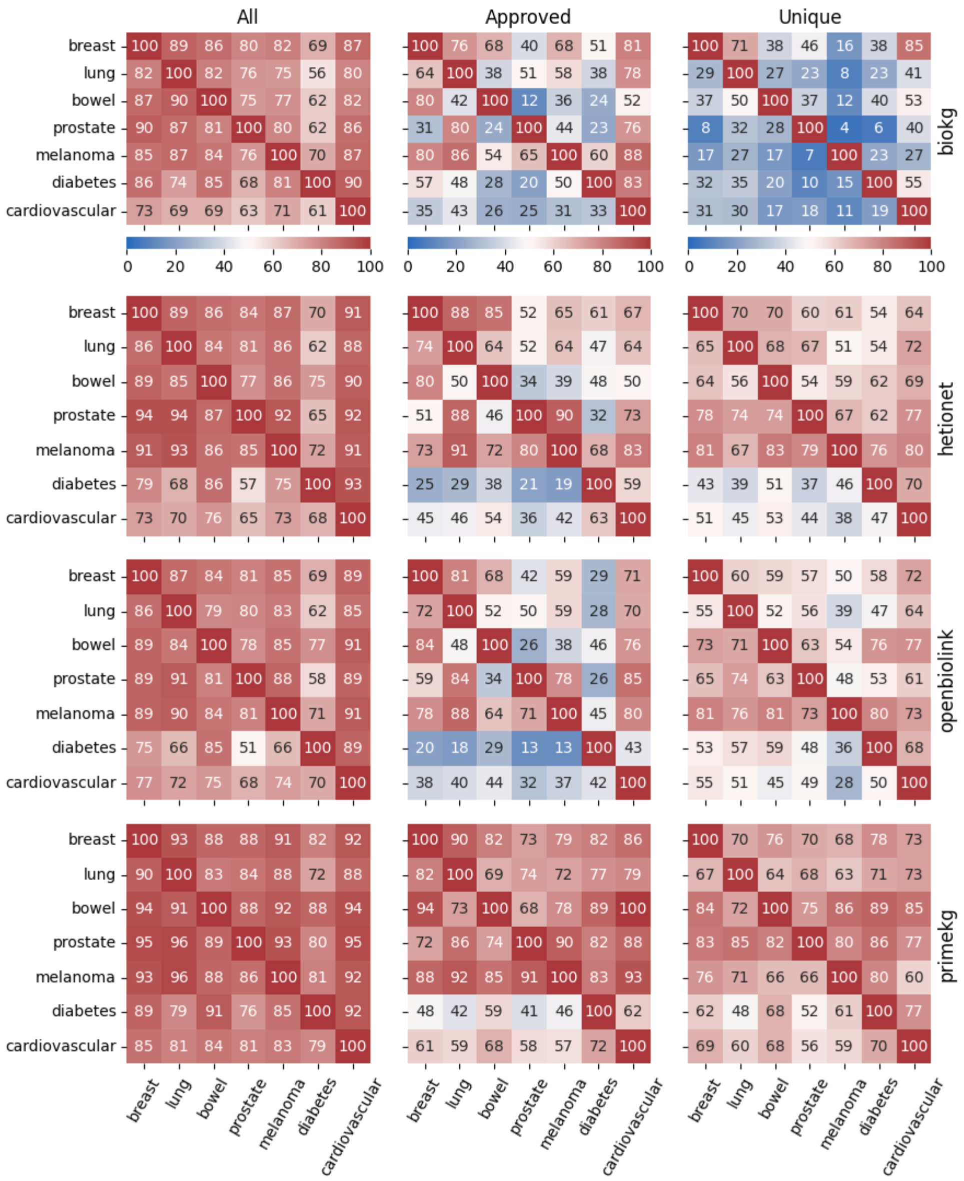


Figure 6. A heatmap showing percentages of common genes between predicted targets from x-axis indications and the input clinical targets (y axis). Each cell is an average of 10 independent results corresponding to 10 different random seeds. No pathway genes were used in training. RF probability threshold of 0.6 was used for model predictions.

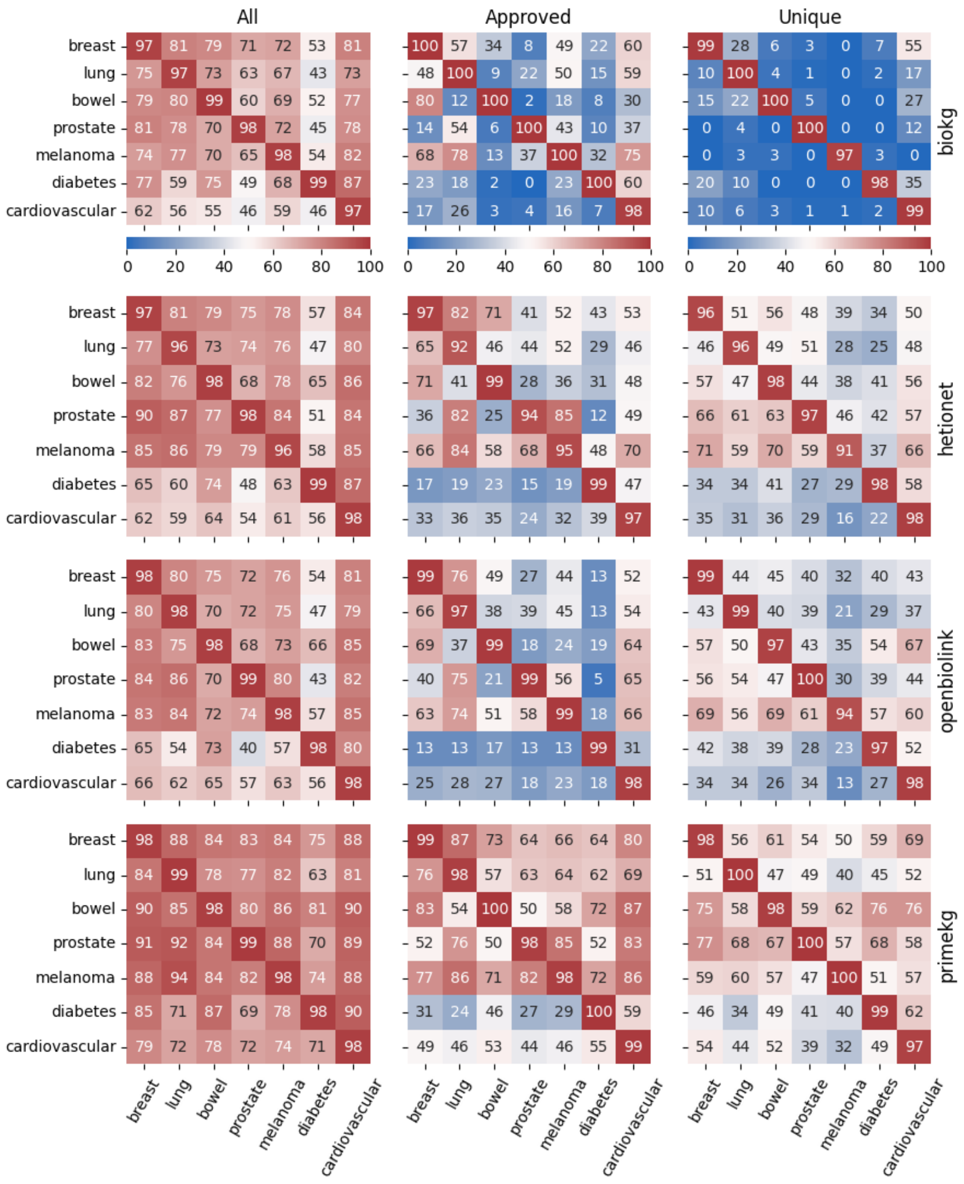


Figure 7. A heatmap showing percentages of common genes between predicted targets from x-axis indications and the input clinical targets (y axis). Each cell is an average of 10 independent results corresponding to 10 different random seeds. No pathway genes were used in training. RF probability threshold of 0.7 was used for model predictions.

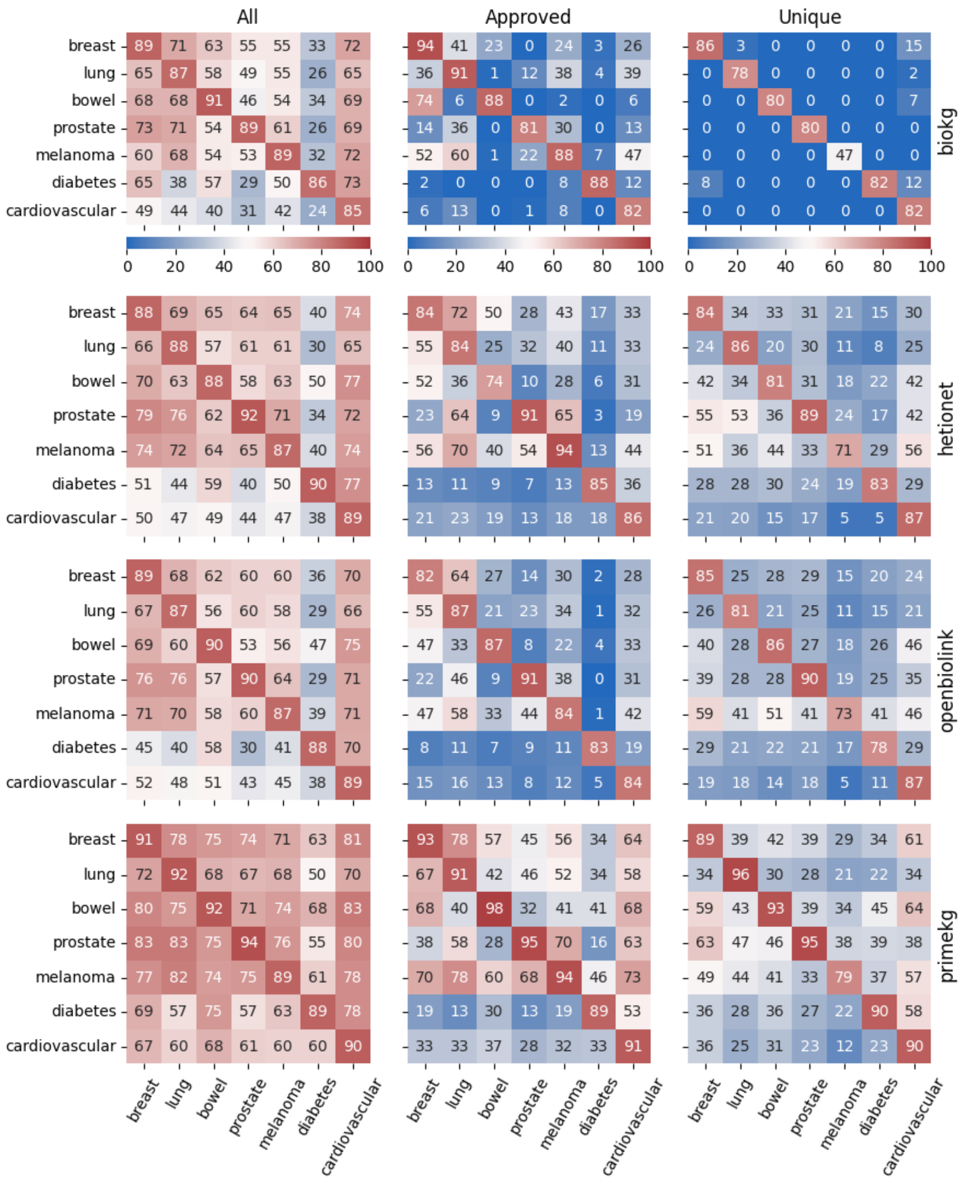


Figure 8. A heatmap showing percentages of common genes between predicted targets from x-axis indications and the input clinical targets (y axis). Each cell is an average of 10 independent results corresponding to 10 different random seeds. No pathway genes were used in training. RF probability threshold of 0.8 was used for model predictions.

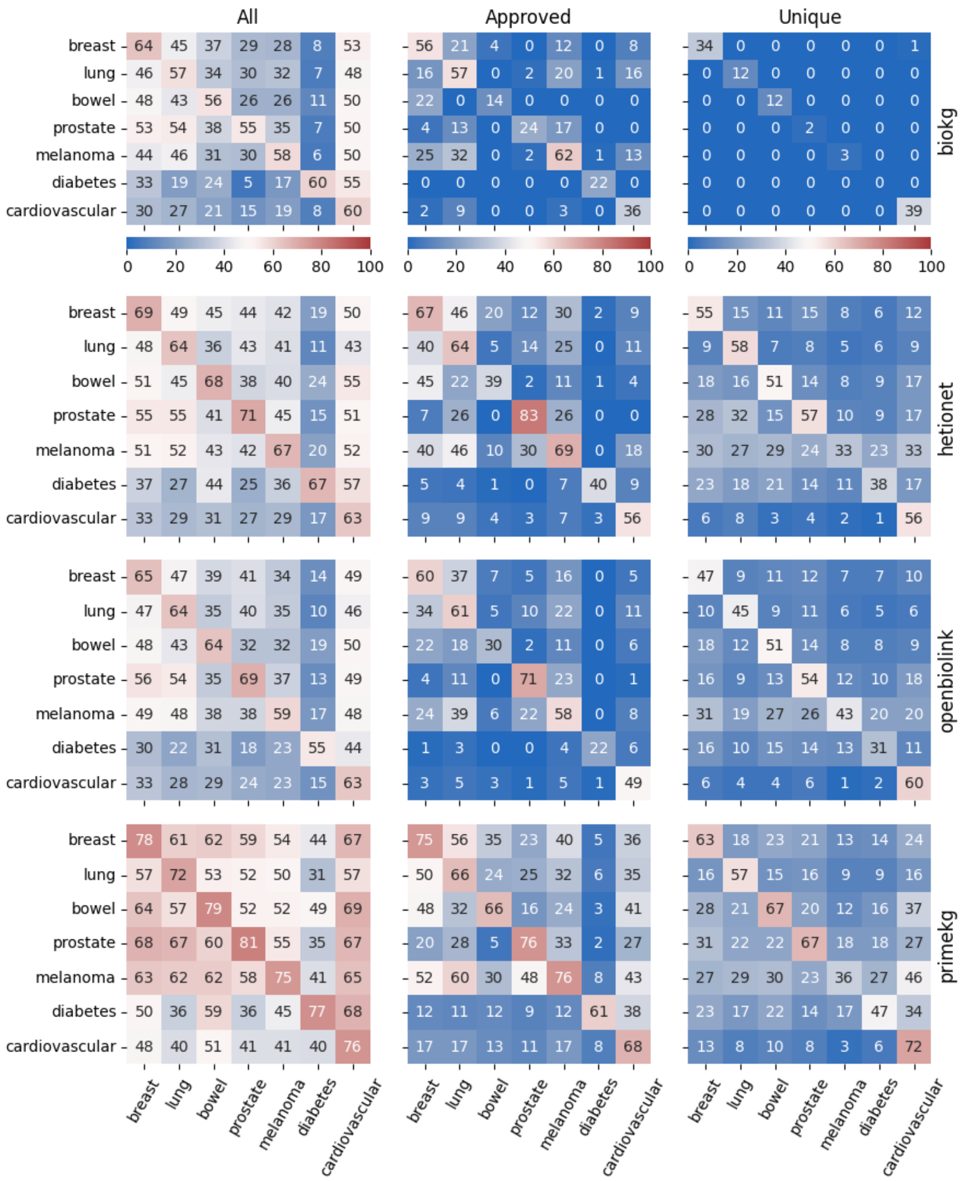


Figure 9. A heatmap showing percentages of common genes between predicted targets from x-axis indications and the input clinical targets (y axis). Each cell is an average of 10 independent results corresponding to 10 different random seeds. No pathway genes were used in training. RF probability threshold of 0.9 was used for model predictions.

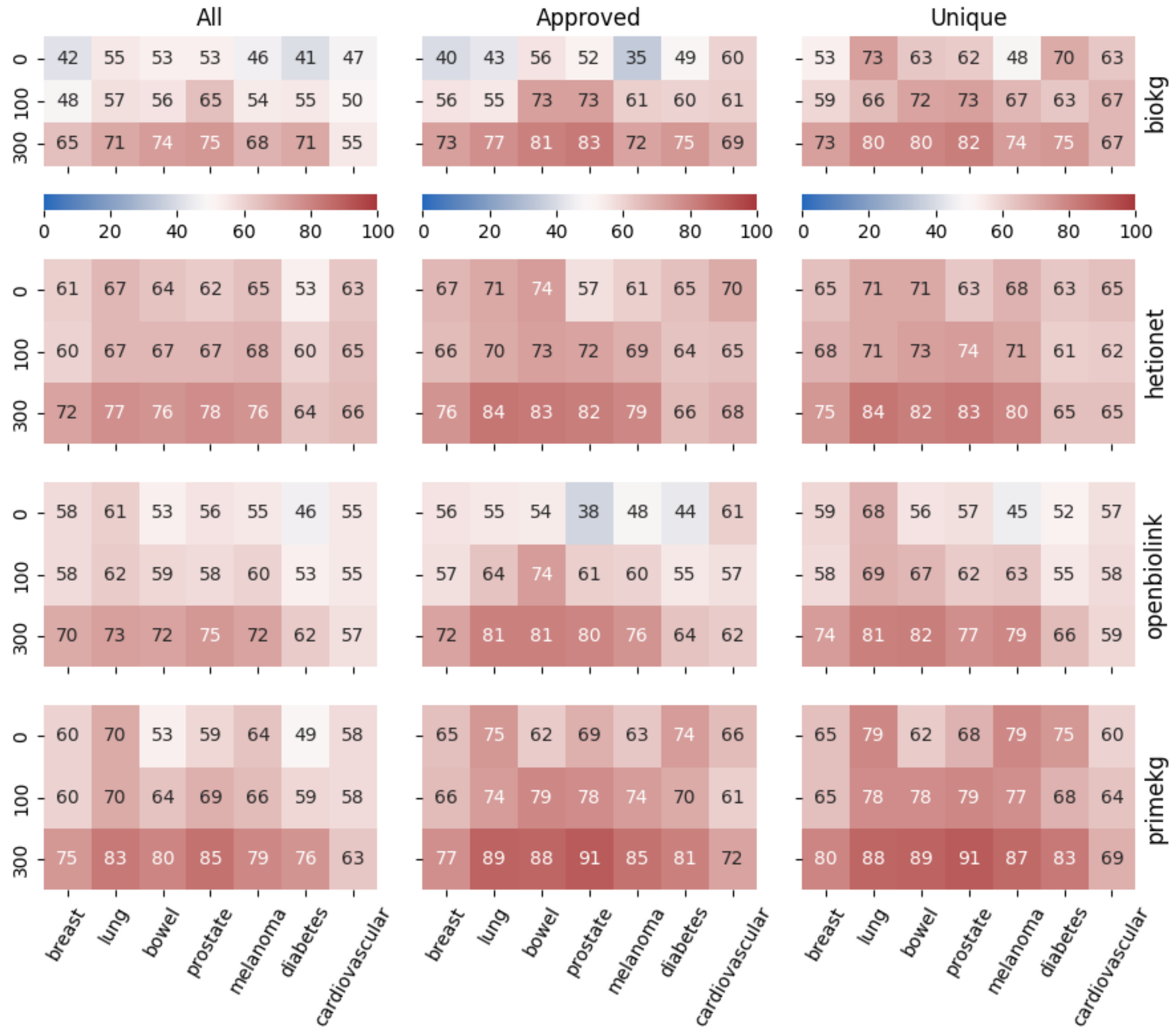


Figure 10. A heatmap (based on 0.5 RF probability threshold) showing percentage of common genes between predicted targets from x-axis indications and the SABCS novel targets. Y axis represents the number of pathway genes used for training (Methods). Each cell is an average of 10 independent results corresponding to 10 different random seeds.

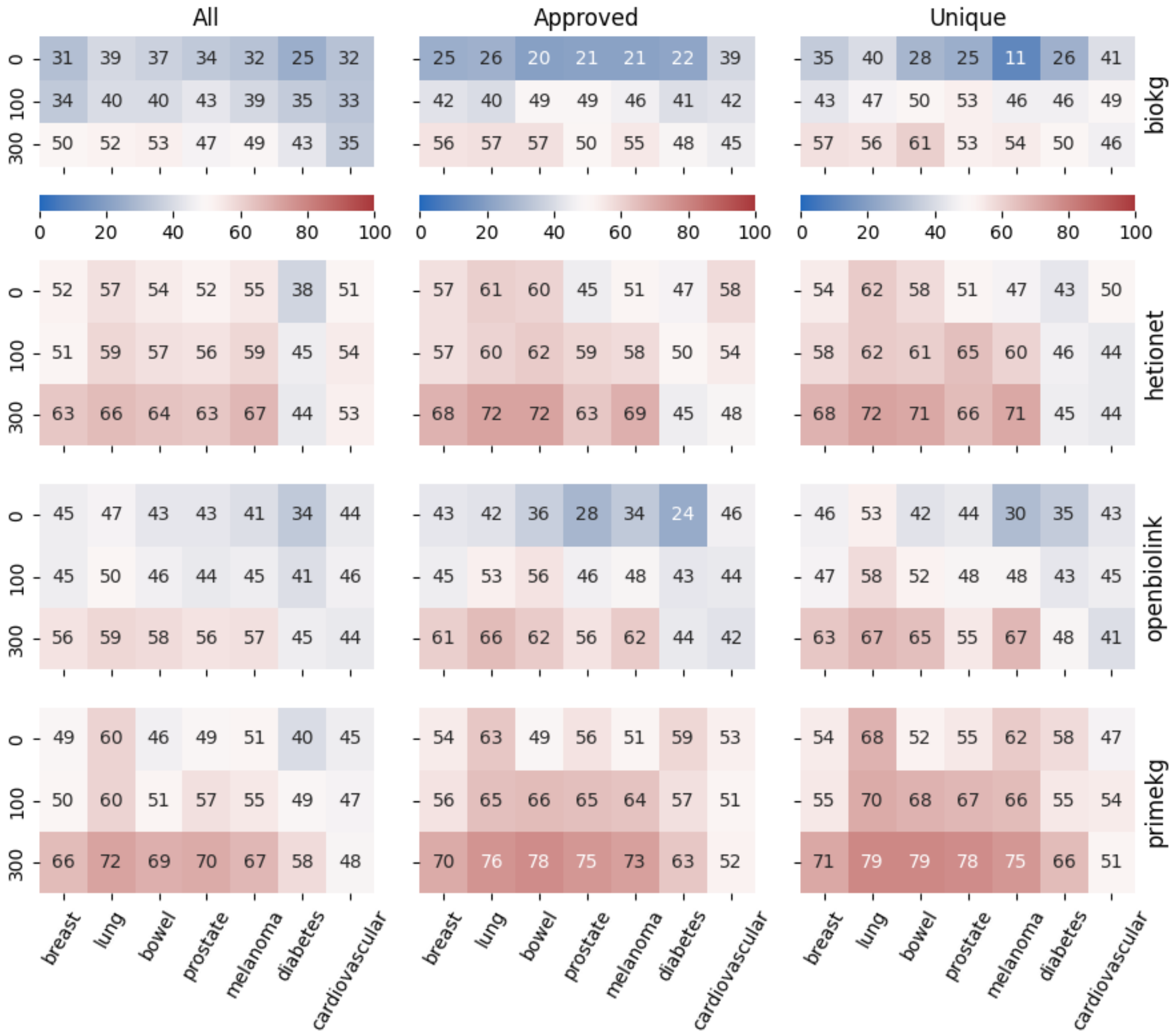


Figure 11. A heatmap (based on 0.6 RF probability threshold) showing percentage of common genes between predicted targets from x-axis indications and the SABCS novel targets. Y axis represents the number of pathway genes used for training (Methods). Each cell is an average of 10 independent results corresponding to 10 different random seeds.

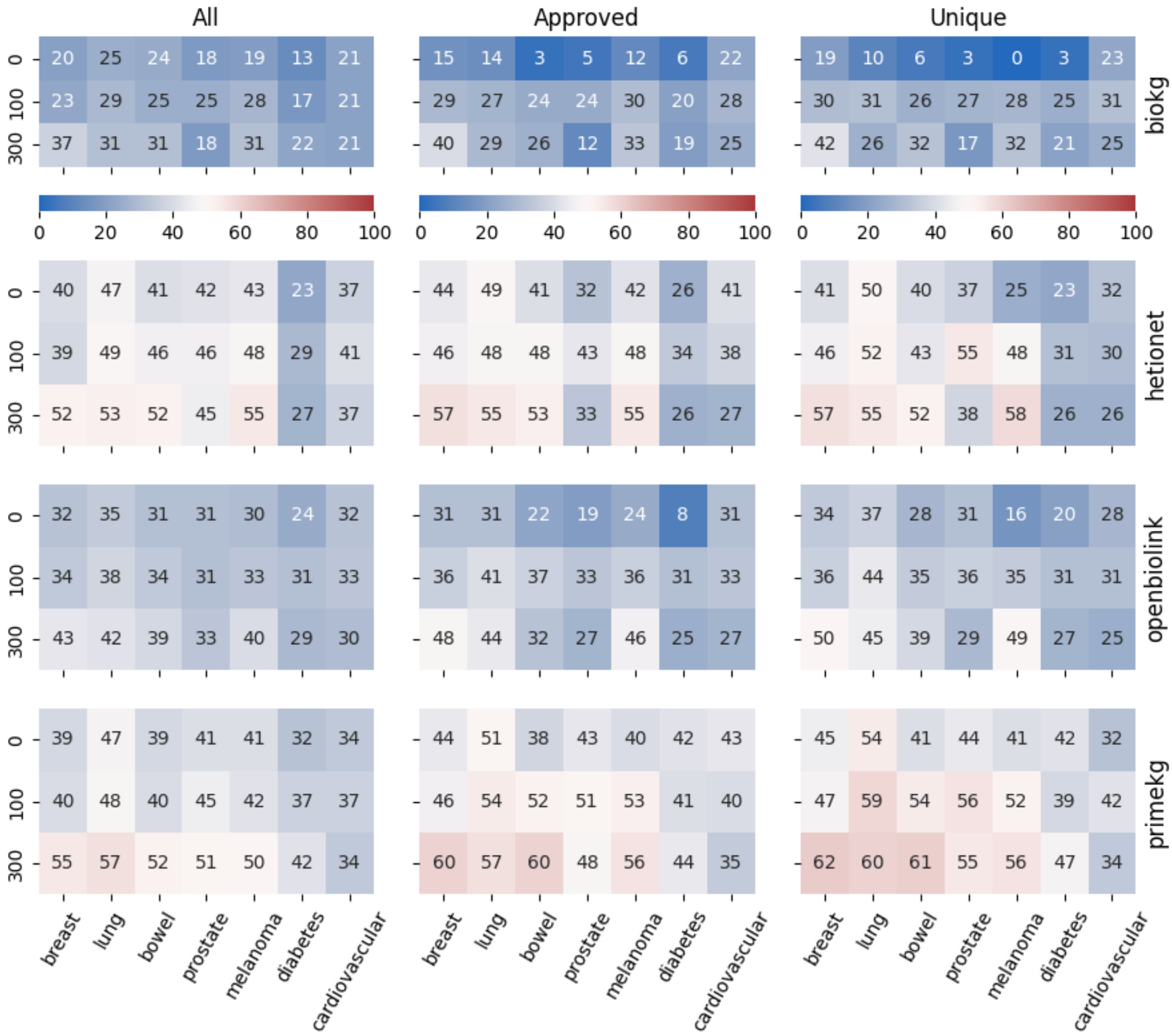


Figure 12. A heatmap (based on 0.7 RF probability threshold) showing percentage of common genes between predicted targets from x-axis indications and the SABCS novel targets. Y axis represents the number of pathway genes used for training (Methods). Each cell is an average of 10 independent results corresponding to 10 different random seeds.

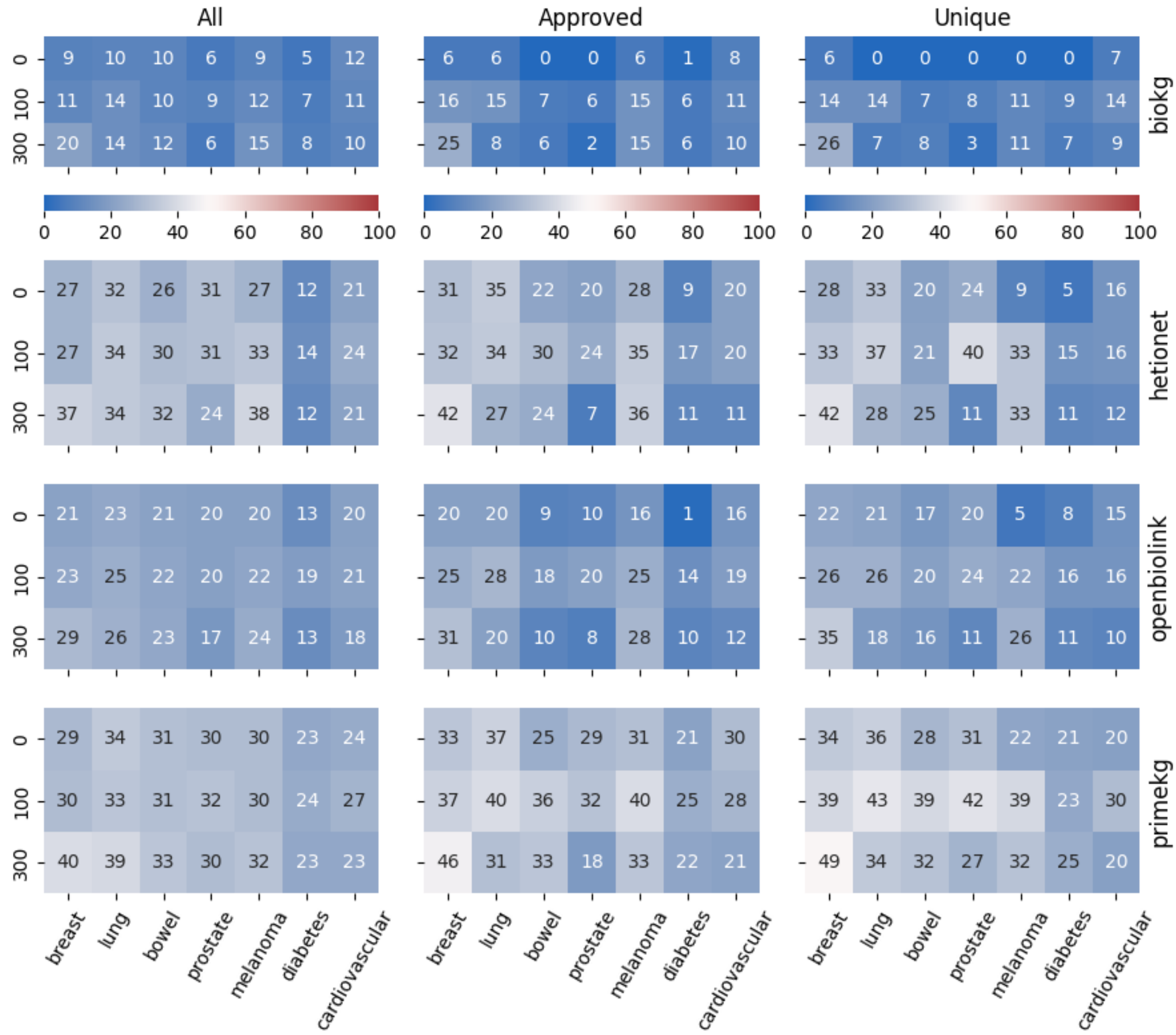


Figure 13. A heatmap (based on 0.8 RF probability threshold) showing percentage of common genes between predicted targets from x-axis indications and the SABCS novel targets. Y axis represents the number of pathway genes used for training (Methods). Each cell is an average of 10 independent results corresponding to 10 different random seeds.

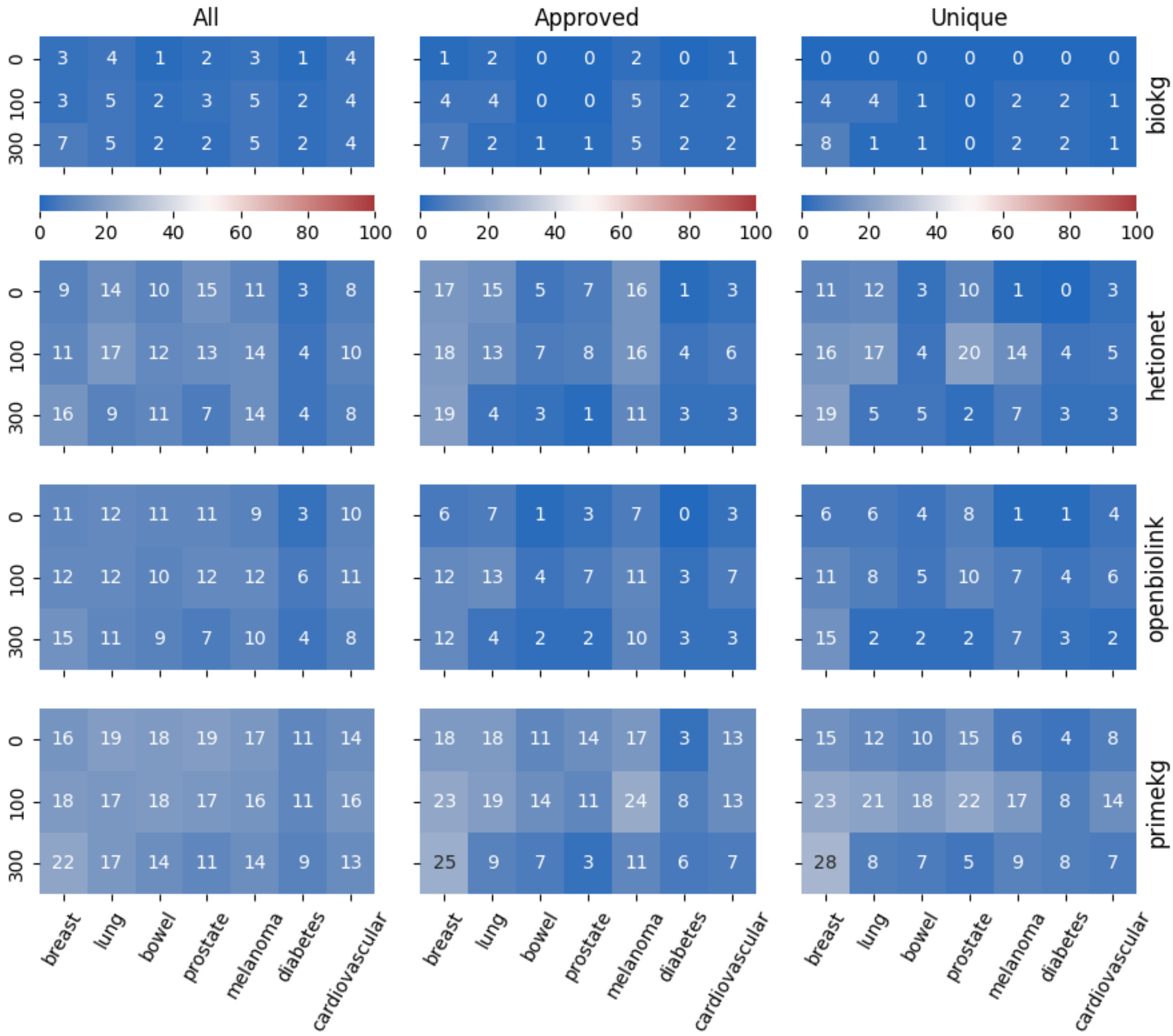


Figure 14. A heatmap (based on 0.9 RF probability threshold) showing percentage of common genes between predicted targets from x-axis indications and the SABCS novel targets. Y axis represents the number of pathway genes used for training (Methods). Each cell is an average of 10 independent results corresponding to 10 different random seeds.
