## Supplementary Notebook for "Artemis: Harnessing Knowledge Graphs for Next-Generation Drug Target Prioritization"

In [1]:
```python
from pykeen.datasets import BioKG, Hetionet, PrimeKG, OpenBioLink
```

### BioKG

#### Summary

In [2]:
```python
BioKG().summarize()
```

```
BioKG (create_inverse_triples=False)
Name        Entities    Relations      Triples
----------  ----------  ------------  ---------
Training    105524      17              1654397
Testing     105524      17               206800
Validation  105524      17               206800
Total       -           -               2067997
Head        Relation                     tail
----------  ---------------------------  -------------
A0A075B6P5  MEMBER_OF_COMPLEX            R-HSA-1479277
A0A075B6P5  PROTEIN_PATHWAY_ASSOCIATION  R-HSA-166663
A0A075B6S6  MEMBER_OF_COMPLEX            R-HSA-1479277
A0A078BQP2  PROTEIN_PATHWAY_ASSOCIATION  R-CEL-2514859
A0A087WPF7  PROTEIN_PATHWAY_ASSOCIATION  R-MMU-8939243
```

#### Relations

All:

In [3]:
```python
list(BioKG().relation_to_id.keys())
```

Out[3]:
```
['COMPLEX_IN_PATHWAY',
 'COMPLEX_TOP_LEVEL_PATHWAY',
 'DDI',
 'DISEASE_GENETIC_DISORDER',
 'DISEASE_PATHWAY_ASSOCIATION',
 'DPI',
 'DRUG_CARRIER',
 'DRUG_DISEASE_ASSOCIATION',
 'DRUG_ENZYME',
 'DRUG_PATHWAY_ASSOCIATION',
 'DRUG_TARGET',
 'DRUG_TRANSPORTER',
 'MEMBER_OF_COMPLEX',
 'PPI',
 'PROTEIN_DISEASE_ASSOCIATION',
 'PROTEIN_PATHWAY_ASSOCIATION',
 'RELATED_GENETIC_DISORDER']
```

Selected protein/gene relations only:

In [4]:
```python
['PROTEIN', 'PPI', 'DPI', 'MEMBER_OF_COMPLEX', 'RELATED_GENETIC_DISORDER'
```

Out[4]:
```
['PROTEIN', 'PPI', 'DPI', 'MEMBER_OF_COMPLEX', 'RELATED_GENETIC_DISORDE
R']
```

### Hetionet

#### Summary

In [5]:
```
Hetionet().summarize()
```

```
Hetionet (create_inverse_triples=False)
Name        Entities    Relations      Triples
----------  ----------  -----------  ---------
Training    45158       24             1800157
Testing     45158       24              225020
Validation  45158       24              225020
Total       -           -              2250197
Head                       Relation    tail
-----------------------  ----------  ------------
Anatomy::UBERON:0000002  AdG         Gene::10005
Anatomy::UBERON:0000002  AdG         Gene::114804
Anatomy::UBERON:0000002  AdG         Gene::118670
Anatomy::UBERON:0000002  AdG         Gene::128989
Anatomy::UBERON:0000002  AdG         Gene::132851
```

#### Relations

All:

In [6]:
```
list(Hetionet().relation_to_id.keys())
```

Out[6]:
```
['AdG',
 'AeG',
 'AuG',
 'CbG',
 'CcSE',
 'CdG',
 'CpD',
 'CrC',
 'CtD',
 'CuG',
 'DaG',
 'DdG',
 'DlA',
 'DpS',
 'DrD',
 'DuG',
 'GcG',
 'GiG',
 'GpBP',
 'GpCC',
 'GpMF',
 'GpPW',
 'Gr>G',
 'PCiC']
```

Selected protein/gene relations only:

In [7]:
```
['GpBP', 'GiG', 'GpPW', 'GpCC', 'GpMF', 'AeG', 'CbG', 'DaG']
```

Out[7]:
```
['GpBP', 'GiG', 'GpPW', 'GpCC', 'GpMF', 'AeG', 'CbG', 'DaG']
```

### OpenBioLink

#### Summary

In [8]:
```
OpenBioLink().summarize()
```

```
You're trying to map triples with 2052 entities and 0 relations that are n
ot in the training set. These triples will be excluded from the mapping.
In total 2047 from 183011 triples were filtered out
You're trying to map triples with 2099 entities and 0 relations that are n
ot in the training set. These triples will be excluded from the mapping.
In total 2093 from 188394 triples were filtered out
```

```
OpenBioLink (create_inverse_triples=False)
Name         Entities    Relations      Triples
----------   ----------  ------------  ---------
Training     180992      28             4192002
Testing      180992      28              180964
Validation   180992      28              186301
Total        -           -              4559267
Head         Relation    tail
----------   ----------  ----------
CL:0000001   IS_A        CL:0000010
CL:0000003   IS_A        CL:0000000
CL:0000006   IS_A        CL:0000101
CL:0000006   IS_A        CL:0000197
CL:0000007   IS_A        CL:0002321
```

#### Relations

All:

In [9]:
```
list(OpenBioLink().relation_to_id.keys())
```

```
You're trying to map triples with 2052 entities and 0 relations that are n
ot in the training set. These triples will be excluded from the mapping.
In total 2047 from 183011 triples were filtered out
```

Out[9]:
```
['DIS_DRUG',
 'DIS_PHENOTYPE',
 'DRUG_ACTIVATION_GENE',
 'DRUG_BINDACT_GENE',
 'DRUG_BINDING_GENE',
 'DRUG_BINDINH_GENE',
 'DRUG_CATALYSIS_GENE',
 'DRUG_INHIBITION_GENE',
 'DRUG_PHENOTYPE',
 'DRUG_REACTION_GENE',
 'GENE_ACTIVATION_GENE',
 'GENE_BINDING_GENE',
 'GENE_CATALYSIS_GENE',
 'GENE_DIS',
 'GENE_DRUG',
 'GENE_EXPRESSED_ANATOMY',
 'GENE_EXPRESSION_GENE',
 'GENE_GENE',
 'GENE_GO',
 'GENE_INHIBITION_GENE',
 'GENE_OVEREXPRESSED_ANATOMY',
 'GENE_PATHWAY',
 'GENE_PHENOTYPE',
 'GENE_PTMOD_GENE',
 'GENE_REACTION_GENE',
 'GENE_UNDEREXPRESSED_ANATOMY',
 'IS_A',
 'PART_OF']
```

Selected protein/gene relations only:

In [10]:
```
['GENE_EXPRESSED_ANATOMY', 'GENE_GO', 'GENE_DRUG', 'GENE_PHENOTYPE', 'GEN
```

Out[10]:
```
['GENE_EXPRESSED_ANATOMY',
 'GENE_GO',
 'GENE_DRUG',
 'GENE_PHENOTYPE',
 'GENE_PATHWAY',
 'GENE_DIS',
 'GENE_GENE']
```

### PrimeKG

#### Summary

In [11]:
```
PrimeKG().summarize()
```

```
PrimeKG (create_inverse_triples=False)
Name        Entities    Relations      Triples
----------  ----------  ------------  ---------
Training    129262      30              6479992
Testing     129262      30               809999
Validation  129262      30               810000
Total       -           -               8099991
Head                                        Relation               tail
-------------------------------------------  ---------------------  ------
---------------------------
'de novo' AMP biosynthetic process           bioprocess_bioprocess  AMP bi
osynthetic process
'de novo' AMP biosynthetic process           bioprocess_protein     ADSL
'de novo' CTP biosynthetic process           bioprocess_bioprocess  CTP bi
osynthetic process
'de novo' GDP-L-fucose biosynthetic process  bioprocess_bioprocess  GDP-L-
fucose biosynthetic process
'de novo' IMP biosynthetic process           bioprocess_bioprocess  IMP bi
osynthetic process
```

#### Relations

All:

In [12]:
```
list(PrimeKG().relation_to_id.keys())
```

Out[12]:
```
['anatomy_anatomy',
 'anatomy_protein_absent',
 'anatomy_protein_present',
 'bioprocess_bioprocess',
 'bioprocess_protein',
 'cellcomp_cellcomp',
 'cellcomp_protein',
 'contraindication',
 'disease_disease',
 'disease_phenotype_negative',
 'disease_phenotype_positive',
 'disease_protein',
 'drug_drug',
 'drug_effect',
 'drug_protein',
 'exposure_bioprocess',
 'exposure_cellcomp',
 'exposure_disease',
 'exposure_exposure',
 'exposure_molfunc',
 'exposure_protein',
 'indication',
 'molfunc_molfunc',
 'molfunc_protein',
 'off-label use',
 'pathway_pathway',
 'pathway_protein',
 'phenotype_phenotype',
 'phenotype_protein',
 'protein_protein']
```

Selected protein/gene relations only:

In [13]:
```
['anatomy_protein_present', 'bioprocess_protein', 'cellcomp_protein', 'di
```

Out[13]:
```
['anatomy_protein_present',
 'bioprocess_protein',
 'cellcomp_protein',
 'disease_protein',
 'drug_protein',
 'exposure_protein',
 'molfunc_protein',
 'pathway_protein',
 'phenotype_protein',
 'protein_protein']
```
